# Decoupling shear stress and pressure effects in the biomechanics of autosomal dominant polycystic kidney disease using a perfused kidney-on-chip

**DOI:** 10.1101/2024.06.18.599137

**Authors:** Brice Lapin, Giacomo Gropplero, Jessica Vandensteen, Manal Mazloum, Frank Bienaimé, Stéphanie Descroix, Sylvie Coscoy

## Abstract

Kidney tubular cells are submitted to two distinct mechanical forces generated by the urine flow: shear stress and hydrostatic pressure. In addition, the mechanical properties of the surrounding extracellular matrix modulate tubule deformation under constraints. These mechanical factors likely play a role in the pathophysiology of kidney diseases as exemplified by autosomal dominant polycystic kidney disease, in which pressure, flow and matrix stiffness have been proposed to modulate the cystic dilation of tubules with *PKD1* mutations. The lack of *in vitro* systems recapitulating the mechanical environment of kidney tubules impedes our ability to dissect the role of these mechanical factors. Here we describe a perfused kidney-on-chip with tunable extracellular matrix mechanical properties and hydrodynamic constraints, that allows a decoupling of shear stress and flow. We used this system to dissect how these mechanical cues affect *Pkd1*^-/-^ tubule dilation. Our results show two distinct mechanisms leading to tubular dilation. For PCT cells (proximal tubule), overproliferation mechanically leads to tubular dilation, regardless of the mechanical context. For mIMCD-3 cells (collecting duct), tube dilation is associated with a squamous cell morphology but not with overproliferation and is highly sensitive to extracellular matrix properties and hydrodynamic constraints. Surprisingly, flow alone suppressed *Pkd1*^-/-^ mIMCD-3 tubule dilation observed in static conditions, while the addition of luminal pressure restored it. Our *in vitro* model emulating nephron geometrical and mechanical organization sheds light on the roles of mechanical constraints in ADPKD and demonstrates the importance of controlling intraluminal pressure in kidney tubule models.

## Introduction

Tubule flow in the kidney results in the application of flow shear stress (FSS), leading to tangential stress on cells; the effect of which has been abundantly described in renal epithelial cell studies^1–4^. The flow rates in the nephron, which decrease in the different segments due to reabsorption events, result in shear stress in the order of 0.2-2 dyn/cm^2^ in renal cells ^1–3^. On renal epithelial cells, flow has been reported to reinforce the apical domain of F-actin cytoskeleton^5^, as well as *adherens* junctions and tight junctions between cells ^2^, to redistribute cytoskeletal tension and remodel focal adhesions ^6^, and to trigger the expression of specific genes, in particular associated with endocytosis and reabsorption^7^. Flow is also intrinsically associated with intraluminal (transmural) pressure, which triggers cell distension^8^ and activates mechanotransduction pathways that are potentially different to those triggered by shear stress. Epithelial renal cells are thus constantly submitted apically to hydrodynamic constraints, flow shear stress, and luminal pressure^9^. In addition, at their basal side they respond to the geometrical and mechanical properties of their underlying basement membrane^10,11^. It is now widely established that cell behavior is deeply influenced by 3D cues. More particularly, the high curvature experienced by cells in the tubule, of typical diameters of 50 µm in the proximal and distal parts, is also an important parameter governing the mechanical responses of renal cells^12,13^

Tubular geometry and mechanical cues are deeply affected in autosomal dominant polycystic kidney disease (ADPKD). In this most common genetic kidney disease (incidence 1/1000-1/2500), mutations (mainly in *PKD1* or *PKD2* genes) inactivating a complex formed by polycystins 1 and 2, result in the development of thousands of kidney cysts in adulthood. This eventually leads to kidney failure, without curative therapies to date so that evolution largely leads to dialysis or transplantation^14^. Moreover, kidney tubules are characterized by their very dense packing, meaning that cyst expansion in ADPKD will lead to mechanical compression of the adjacent tubules. The nephron obstructions due to cyst formation result in altered flow shear stresses and luminal pressures in affected nephrons and their neighbors, which may account for the clustered and exponential propagation of cysts in the disease known as the snowball effect ^15^.

Polycystins 1 and 2 are respectively a large orphan receptor and a Transient Receptor Potential cation channel, coupled to numerous signaling pathways including GPCRs, mTOR, JAK-STAT or Wnt pathways^16,17^. Although the mechanism underlying the initial local tube dilation at the origin of cyst formation is currently thought to rely on perturbations in proliferation, apoptosis and/or planar cell polarity ^18–21^, it remains to date poorly understood. Many aspects, however, point to an important contribution of mechanical factors in cystogenesis. Cells integrate the mechanical stimuli through several organelles, including primary cilia and cell-substrate and cell-cell adhesions, with mechanosensitive properties deeply perturbed in ADPKD^10,22,23^. Polycystins are present in the primary cilia, a sensor of the luminal flow, although their mechanisms of action remain elusive^24,25^. However, primary cilia ablation reverts the cystic phenotype in ADPKD, pointing to the primary involvement of this organelle in the disease^26,27^. Moreover, transcriptomic assays comparing the effect of FSS on WT or *Pkd1^-/-^* proximal cells found that several genes were slightly more induced in the latter condition, suggesting that *Pkd1* here has a role in restraining FSS-regulated signaling ^28^.

In addition, polycystins intervene in the organization of integrin-based basal complexes as well, with overexpression of several integrins and their extracellular matrix ligands in ADPKD ^29^, along with a reversion of cystic phenotypes reported upon β1 integrin inactivation ^30^. Polycystins are also strongly involved in acto-myosin organization ^10^ and actin-based mechanotransduction ^31^, with RhoA-YAP-c-Myc axis identified as a key player in cystogenesis ^32^ that participates in stiffness sensing ^10^.

Studying biomechanical aspects of cyst formation in a mice model of ADPKD, we recently discovered that *Pkd1* loss triggers the remodeling of the distal nephron basement membrane, leading to tubule distension, tubular cell stretching and increased tubule distensibility. In this context, ureteral obstruction, which increases intraluminal pressure resulted in an excessive dilation of the distal nephron followed by explosive cystogenesis suggesting that increased luminal pressure could be a biomechanical driver of ADPKD^33^. Few studies have addressed the specific effects of pressure dependence on polycystin expression or function^34–37^, including mechanisms on renal cells relying on the inhibition of Piezo1-dependent stretch-activated channels^35^ by polycystin-2. But this electrophysiology work was not performed in the frame of a realistic tubule geometry. The influence of decoupling these contributions for tube dilation and cyst formation in ADPKD remains unknown, despite the importance of identifying their respective role in the elucidation of the mechanosensitive pathways involved and for future therapeutic applications.

A variety of 3D models reproducing realistic features of the kidney tubules, as well as some pertinent mechanical properties like perfusion or substrate elasticity, have been developed in the past few years. These models involve organoids ^38,39^, bioprinting tubules ^40,41^, or organs-on-chips based on wire molding techniques ^42^. However, the coupling of flow shear stress with hydraulic pressure in most experimental devices results in the fact that not only the relative contributions are not distinguished, but also that some effects observed can be attributed misleadingly to shear stress alone. In *in vitro* tubes, FSS and pressure contributions can however be decoupled by playing on the hydrodynamic properties of the system, like hydraulic resistance, diameter, or material mechanical properties, as described recently for endothelial cells ^43^.

In our past works, we developed a kidney-on-chip specifically dedicated to the study of ADPKD, by reproducing tightly packed tubes of physiological diameter in a biocompatible deformable collagen substrate, in which we observed a tube dilation specific for ADPKD proximal cells^44^ . We recently implemented versatile possibilities of perfusion on our devices ^45^. In this paper, we develop an original system to decouple flow and luminal pressure in our kidney-on-chips. This allows us to dissect the effects of flow alone, flow with luminal pressure, and matrix stiffness on tubule dilation, proliferation, and multicellular morphology. We find contrasting mechanisms of tube dilation, driven or not by proliferation, depending on cell lines that originate either from proximal or distal parts of the nephron. Importantly, our results highlight the prominent role of luminal pressure for tube dilation in collecting duct cells, which is of paramount importance as cysts form first in this nephron part in ADPKD.

## Methods

### Kidney-on-chips

As previously described ^45^, the chip consisted of a PDMS scaffold of 3 PDMS layers bonded to a glass slide. One part of the scaffold featured the microfluidic channels. For this part, a PDMS (Curing Agent to PDMS weight ratio of 1:10) was poured on a mold obtained with photolitography. This part laid on a flat PDMS membrane which elevated the channels from the glass slide. This flat PDMS layer was obtained by spincoating PDMS 1:10. On top, another flat 500 µm thick PDMS membrane closed the chip. The PDMS scaffold surface was activated by oxygen plasma and then treated with a 5% (3-aminopropyl) triethoxysilane (Sigma-Aldrich) solution in methanol for 45 min, rinsed thoroughly in methanol in an ultrasonic bath then in a 2.5% glutaraldehyde (Sigma-Aldrich) solution in ddH20 for 45min and rinsed thoroughly in ddH20 in an ultrasonic bath then at room temperature in ddh20 overnight. Then, 75 µm wide tungsten wires were inserted into the scaffold, which was sterilized with UV-Ozone. A 6mg /mL or 9 mg/mL collagen I gel was prepared on ice by neutralizing high concentration collagen gel from rat tail with NaOH and diluted in PBS with 10% of total volume with PBS 10X. The gel was injected in the collagen chambers, still on ice. The collagen was set at 37°C for 2h. Chips were then stored in PBS at 4°C up to 2 weeks before use.

### Flow rate measurements

5 µm fluorescent beads (Polysciences) in PBS were used. The flow of beads was imaged with a fluorescence microscope, and bead tracking was performed with ImageJ and the extension Trackmate. From this data, the average flow speed of beads within a channel was calculated. The flow rate corresponding to the channel was then computed. We then estimated the wall shear stress in an 80µm micron wide tubule with the same flow rate.

### Cells

Mouse PCT *Pkd1^+/-^* and *Pkd1^-/-^* cells (respectively PH2 and PN24 clones) were a kind gift of S. Somlo ^46,47^. These cells, containing the Immortomouse transgene for the interferon-inducible expression of a thermolabile large tumor antigen, were amplified in proliferation conditions (33°C, with γ-interferon) and differentiated in differentiation conditions (37°C, without γ-interferon). Proliferation conditions were 33°C, 5% CO2, in DMEM/HamF12 supplemented with 3% SVF, 7.5 nM Na selenite, 1.9 nM triiodo-L-thyronine, 5 mg/ml insulin, 5 mg/ml transferrin, 100 UI/ml penicillin/streptomycin, 5 mg/ml nystatin (all from Sigma), and 10UI/ml γ-interferon (Millipore). Cells were differentiated in the same media without γ-interferon, and with 1% SVF instead of 3% SVF, at 37°C, 5% CO2.

Edited mIMCD-3 cell line (immortalized cells from murine inner medullary collecting duct, deleted for *Pkd1* gene using CRISPR-based genome editing or corresponding control (parental), were kindly provided by Michael Köttgen^48^ and cultivated in DMEM/HamF12 supplemented with 10% SVF at 37°C, 5% CO2.

### Tube seeding and perfusion

The day before seeding, wires were partially removed to form the channels, an anti-adhesion solution was perfused in the tubes during a few seconds followed by a 1-hour incubation at room temperature, and a laminin 50µg/mL solution was injected in the chip using a pressure controller and PTFE tubing followed by 1 hour-incubation at 37°C. Cells were detached from flask upon confluency with trypsin, counted and resuspended at 10 million cells /ml in complete medium. The cell suspension was then injected into collagen channels with a pressure controller via PTFE tubing until the channels were filled with cells. Then tungsten wires were inserted back into the channels to form a hollow tube. Chips were incubated in culture medium for 2 to 3 hours to let cells adhere, then tungsten wires were removed. Before perfusion, the chip channel was flushed with a pressure controller (Fluigent Flow E/Z) at 50 mbar with a 37°C PBS solution to remove unattached cells and put back in the culture medium.

Chips were perfused with 3 ml TERUMO syringes attached to a microfluidic needle and maintained vertically by a homemade holder. Chips were then cultured in 6-well plates in humid atmosphere at 37°C for up to 1 week, with medium changed every two days.

### Tube labeling

Chips were washed with PBS twice before being fixed in a 4% paraformaldehyde solution in PBS for 45 min. Chips were then rinsed twice in PBS and stored in PBS at 4°C before immunostaining according to our previous described procedure ^45^. Cells were permeabilized by submerging the chip into PBS + 0.3% TritonX100 for 30 minutes and rinsed three times for 30 minutes with PBS-BSA 2%. The chip was submerged in PBS-BSA 2% with 0.1% Tween20 and 2% Goat Serum for saturation. A 100-150 µL drop of 1/200 antibody, 2% BSA, 1% Tween20 in PBS is delicately placed on the chip so that it does not fall. Incubation is carried out for 24 hours in a humid environment at room temperature. The chip was rinsed 3 times for 30 minutes in PBS-BSA 2%. A 100 µL drop of a solution containing 1 µg/ml Hoechst, 0.25 µg/ml fluorescent phalloidin, primary antibody (rabbit anti-Ki67, Abcam, ab16667, 1:100), 2% BSA, 1% Tween-20 in PBS was delicately placed to prevent spilling. Incubation was carried out for 24 hours in a humid environment at room temperature. The secondary antibody was a 641 nm anti-rabbit highly cross-adsorbed antibody. Imaging was performed by confocal microscopy.

### Quantifications

Measurement and normalization of tube diameter was performed as previously described^44^. Nuclei segmentation and identification of Ki67 positive nuclei was performed in two steps, first presegmentation with the software Ilastik and construction of a training pool with each pixel assigned to nuclei, actin, or background, second segmentation, with a home-made Matlab algorithm. To compute the internuclear distance, for each nucleus, the 6 closest nuclei were determined. To filter out outliers due to border localization or isolated nuclei, we considered that the distances between nuclei and their 6 closest neighbors should follow a gaussian distribution, and only elements falling between the 1st and 3rd quartile of such distribution was kept. Then distances between nuclei and their 6 closest neighbors were averaged over each nucleus, and finally averaged over all the nuclei. To compute Ki67 positive ratio, Ki67 positive nuclei were automatically identified and counted. Then total number of Ki67 cells were divided by total number of nuclei.

### Statistical analysis

All statistical analyses were performed using the software Prism. Tubular dilation was analyzed with two tailed t-tests. Ki67 positive cell ratio were analyzed with separate two-tailed t-test between cell type for each day. Internuclear distances were analyzed with two-way ANOVA.

### COMSOL simulation

We used the COMSOL Multiphysics® software (version 5.5, COMSOL AB, Stockholm, Sweden) to model fluid flow and luminal pressure in the kidney-on-chip. The simulations were conducted using a 3D model, which included the 5 rectangular serpentine channels followed by the round collagen channels. Each serpentine has a section with a width and height of 50 µm and were 6cm long in total. Each collagen channel had an 80µm diameter. A creeping flow was set along the channels with pressures in inlet 1 and 2 as boundary conditions. The inlet 1 pressure was set to 10 mbar in the flow configuration and 0 mbar in the flow and pressure configuration. The inlet 2 pressure was set to 0 mbar in the flow configuration and 10 mbar in the flow and pressure configuration. The computational mesh employed was the default fine physics-controlled mesh provided by COMSOL. For pressure, data were visualized with slice plots in the mid plan of the chip, extending to the total length of the microfluidic channels. For shear stress, data were visualized with slice plots showing the mid circular cross section of the collagen tubules.

## Results

### 1. Decoupling flow and luminal pressure in kidney-on-chips

To finely dissect the roles of flow shear stress and intraluminal pressure within the kidney tubule independently, we first optimized our previously developed kidney-on-chip system^45^ (Figure 1.A). Briefly, a gel of collagen type I was polymerized in a polydimethylsiloxane (PDMS) microfluidic chip; this gel featured parallel 80µm hollow tubules where kidney cells of two mouse cell lines were seeded, PCT and mIMCD-3 cells. The chip not only integrated these collagen tubules but also highly resistive serpentine microfluidic channels in the PDMS bulk perfusing the tubules, which allowed to decouple flow shear stress from intraluminal pressure. Flow and pressure in chips were generated by a home-made system, consisting in medium-filled syringes controlling the hydrostatic pressure; these simple devices allowed to perform the experiments in many chips in parallel. The chips could be perfused using two different inlets (Figure 1.A-B) and, depending on the inlet used and due to the important pressure drop downstream the serpentines, the intraluminal pressure in tubules was either negligible (inlet 1) or equivalent to the set inlet pressure (inlet 2). In both cases, the resulting flow shear stress would then in theory be the same, as was verified by a finite element simulation (Figure 1.C). We chose to perfuse the chip with an input of 10 mbar pressure as a typical intraluminal pressure in the kidney nephron. The serpentine geometry was set so that this pressure generates a flow shear stress in the tubules in the 0.1-1 dyn/cm², as confirmed by a finite element simulation (Figure 1.C).

**Figure 1:**
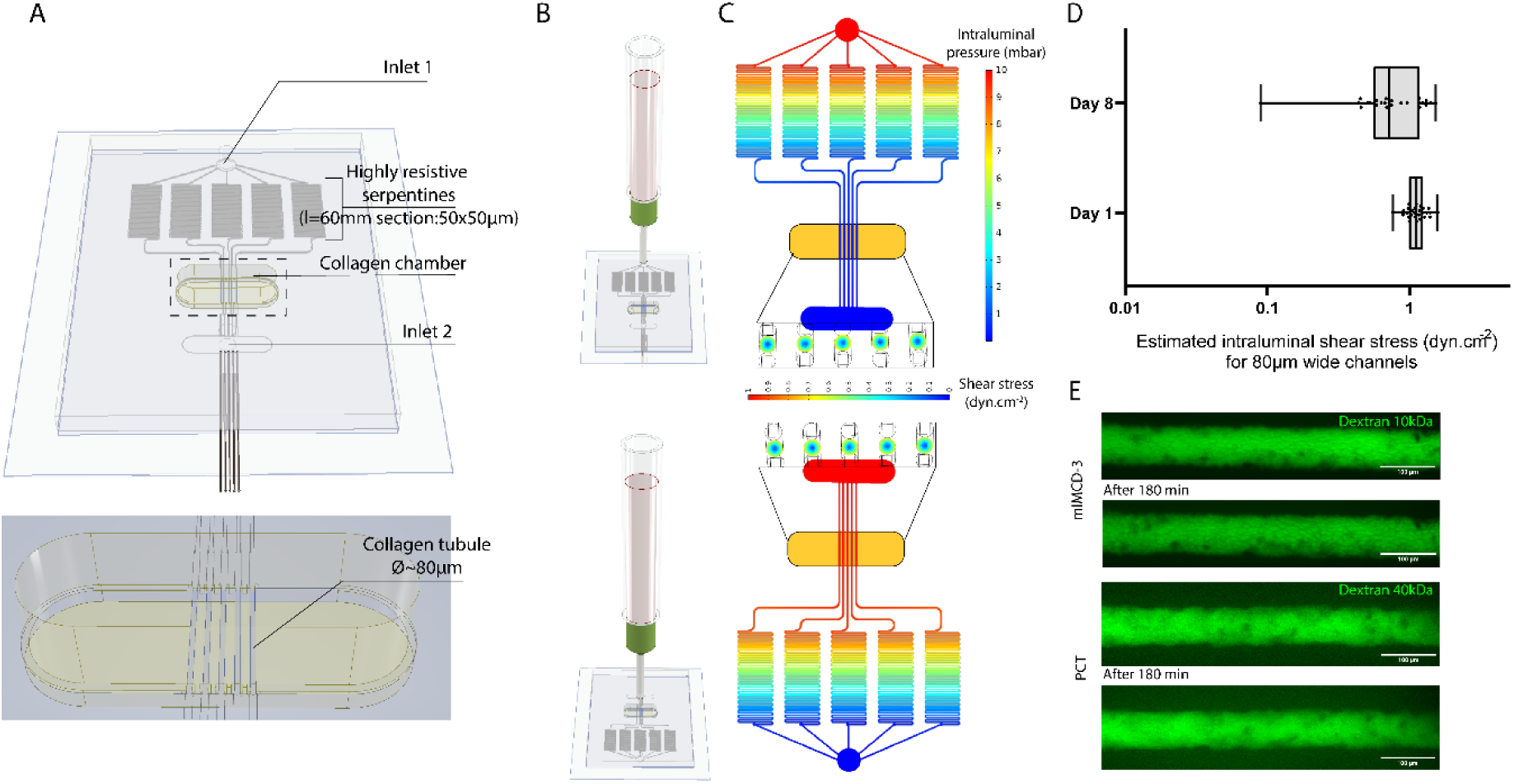
A perfused kidney-on chip to decouple flow shear stress from intraluminal pressure. A 3D model of the chip pointing at its key features is shown in **(A)** with a zoom on the collagen microtubules. Depending on where the chip is perfused **(B)**, either in the inlet 1 (top) or in the inlet 2 (bottom), we show with a finite element simulation **(C)** (COMSOL Multiphysics) that a pressure of 10 mbar will result in a negligible intraluminal pressure or in a 10-mbar intraluminal pressure respectively with the same flow shear stress in the tubules around 1 dyn/cm^2^. **(D)** Estimation of the intraluminal shear stress for 80µm wide tubules, calculated from measurements of the flow rates in confluent tubules of mIMCD-3 cells 1 day (n=27) and 8 days (n=20) after seeding. (Boxplots showing are 1^st^ quartile, median and 3rd quartile, with whiskers showing extrema. Each dot represents a tubule.) **(E)** Impermeability of m-IMCD-3 (top) and PCT (bottom) tubules to FITC Dextran 10 kDa and 40 kDa respectively. Pictures show the temporal evolutions of one mIMCD-3 and one PCT tubule at the initial time point and after 180 min.

We experimentally measured the flow rates in tubules seeded with mIMCD-3 cells by particle tracking with 5µm fluorescent beads, to verify if the system was indeed in the expected range (Figure S1.A). From these measurements, we estimated the flow shear stress on cells lining an 80µm wide channel. We observed that 1 to 8 days after seeding, this flow shear stress remained within physiological range (Figure 1.D), although there was a notable drop in flow shear stress certainly due to cell clogging. For this reason, we restrained our study to 8 days after cell seeding. We also verified the tightness of the tubules obtained from mIMCD-3 cells and PCT cells. After perfusing confluent tubules with fluorescent dextran, looking at the fluorescence of the medium in and around the tubules, we demonstrated that mIMCD-3 cells formed a barrier for diffusion of molecules as small as 10 kDa. PCT tubules on the other hand appeared leaky for 10kDa dextran, but impermeable to 40 kDa dextran (Figure 1.E). This confirmed that the expected intraluminal pressure could build up within the tubules. Indeed, a perfusion in the flow and pressure configuration tubules lined with a confluent layer of kidney cells almost immediately led to a slight tubular dilation, not observed with naked collagen tubules (Figure S1.B).

### 2. Hydrodynamic constraints differentially affect the ability of Pkd1-/- mIMCD-3 and PTC cells to dilate kidney tubule-on-chip

We used cell lines generated from the proximal tubule (*Pkd1^+/-^* PCT cells) and from the collecting tubule (*Pkd1^+/+^* mIMCD-3 cells), from which *Pkd1*^-/-^ cells were generated. We previously reported the behavior of these cells in a similar tubular microenvironment but in static conditions^44^, where both *Pkd1*^-/-^ cell lines dilated the tubules significantly more than their parental cells (Myram et al. 2021^44^, Mazloum et al.^33^, Figure S2). After validation of our perfused system, we investigated the behavior of these cells submitted to a 1 dyn/cm² flow shear stress, combined or not with a 10- mbar intraluminal pressure. Parental PCT cells did not dilate tubules when subjected to flow shear stress only (Figure 2.A), while a significant dilation of *Pkd1*^-/-^ tubules was observed 5 days after confluency, between 30 and 45% (95%CI). Interestingly, with flow shear stress only, the loss of *Pkd1* in mIMCD-3 cells was no longer sufficient to trigger significant tubular dilation (Figure 2.B). Combining FSS with a 10-mbar intraluminal pressure led to rapid tubular dilation of both cell types after 1 day. At this early time, there was no significant difference between genotypes, with tubular diameter increasing by 26-40% and 32-58% for PCT parental and *Pkd1*^-/-^ cells respectively (Figure S3.A), and by 70-92% and 70-99% for parental and *Pkd1*^-/-^ mIMCD-3 cells (Figure S3.B). Nevertheless, within 5 days, total dilation was clearly enhanced by the loss of *Pkd1*, with PCT tubules dilating by 49-75% for parental cells but 86-125% for *Pkd1*^-/-^ cells (Figure 2.C, left). The combination of FSS with intraluminal pressure restored the excessive dilation of mIMCD-3 *Pkd1*^-/-^ tubules, with dilation levels reaching 2.3-2.9 times (95% CI) - almost 4 times in extreme cases - the initial diameter after 5 days (Figure 2.D, left). These results suggested that dilation specific to the *Pkd1* status occurred mostly between day 1 and day 5. Worthy of note, we also observed a different pattern between PCT and mIMCD-3 parental cells. If parental PCT tubules still dilated by 20% on average past the initial dilation, there was only little deformation of parental mIMCD-3 tubules with dilation levels after D1 well below 10% (Figure 2.C-D right, D5/D1).

**Figure 2:**
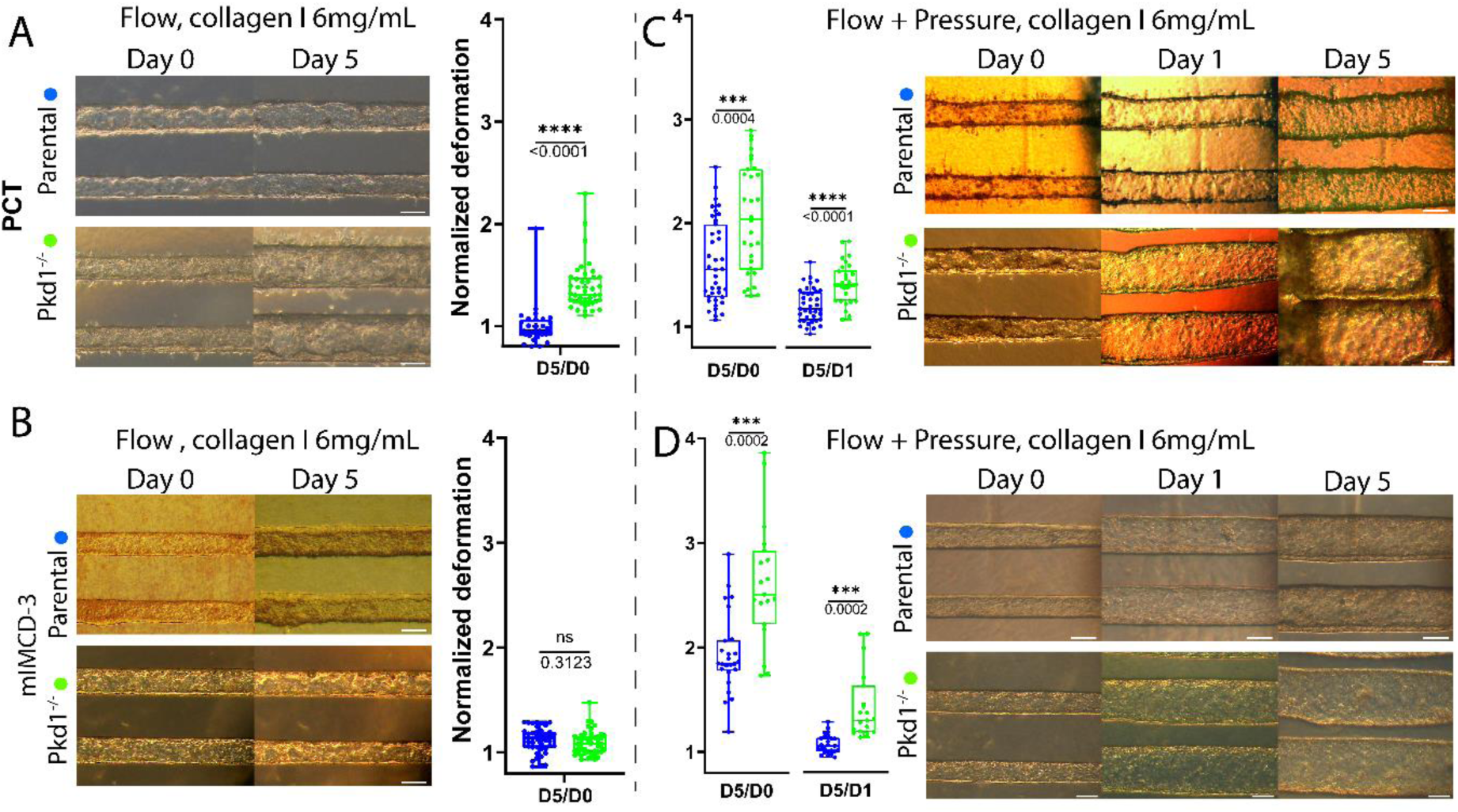
Under flow shear stress, loss of *Pkd1* leads to increased enlargement of PCT tubules, but intraluminal pressure is necessary for increased enlargement of mIMCD-3 tubules. PCT **(A,C)** or mIMCD-3 **(B, D)** tubules were perfused after seeding without intraluminal pressure (A,B) or with intraluminal pressure (C,D). Representative tubules are shown in these configurations, on the day of confluency and one or several days after confluency. Mean diameters of the tubules 5 days after confluency normalized by their mean diameters at the day of confluency (D5/D0) are indicated. In the flow+pressure condition, characterized by *Pkd1^-/-^* dilation for both cell types, the evolution following the initial dilation is also quantified, with diameters at day 5 normalized by diameter at day 1 (D5/D1, right). Parental PCT cells (blue, A, flow n=29, C, flow+pressure n=39), PCT *Pkd1^-/-^* cells (green, A, flow n=41, C, flow+pressure n=30), parental mIMCD-3 cells (blue, B, flow n=41, D, flow+pressure n=25) and mIMCD-3 *Pkd1^-/-^* cells (green, B, flow n=47, D, flow+pressure n=19). Each dot represents a tubule (ns: p-value>0.05, *: p-value≤0.05, **: p-value≤0.01, ****: p- value≤0.0001, actual p-values are indicated below the symbols). Scale bars: 100µm.

### 3. Pkd1-dependent tube dilation upon flow + pressure is prevented by increasing the collagen concentration for mIMCD-3 cells, but not for PCT cells

As seen above, an intraluminal pressure of 10 mbar combined with FSS was sufficient to trigger tubular dilations, regardless of *Pkd1* status or cell type, in a 6mg/mL collagen I matrix. As *ex vivo* perfusion experiments demonstrated^49^, the matrix surrounding kidney tubules, more specifically the basement membrane, constitutes the principal barrier preventing tubular dilation when intraluminal pressure increases. To assess the importance of the surrounding matrix properties in *Pkd1*-dependent tubular dilation mechanisms, we cultivated on-chip mIMCD-3 and PCT cells in scaffolds with increased collagen I concentration (9 mg/mL; Figure 3). We previously measured by tensile tests the Young modulus of 6 and 10 mg/mL collagen I gels employed in our study at 55±15 kPa to 86±47 kPa respectively (Bernheim-Dennery et al., unpublished data). The relative increase in Young modulus thus remains moderate and is corroborated by a comprehensive review of existing literature, where we observe a quadratic relationship between the gel’s Young’s modulus and its concentration^50–53^.

**Figure 3:**
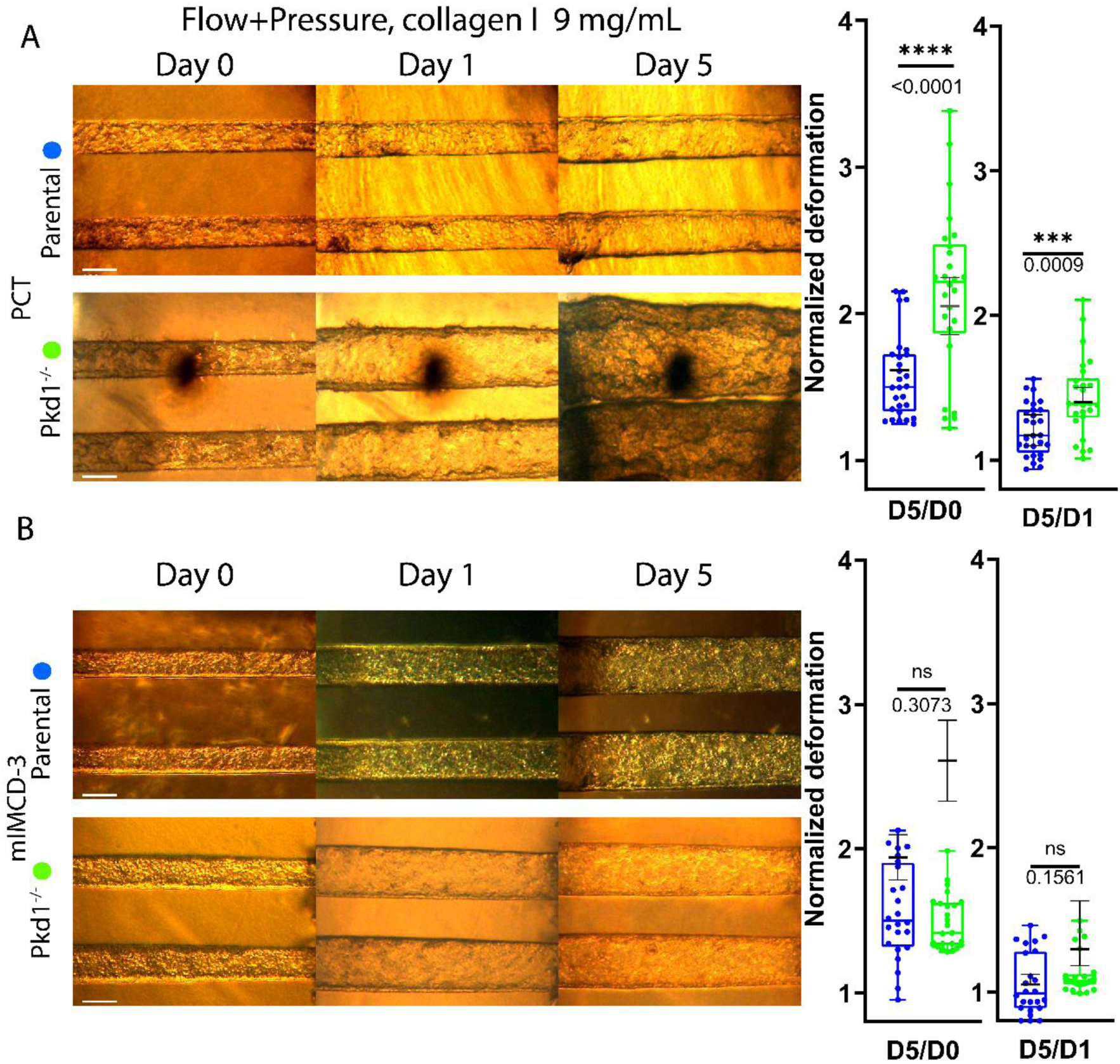
Increase in pressure-driven tubular dilation due to loss of *Pkd1* is mitigated by denser extracellular matrix in mIMCD-3 tubules but not in PCT tubules. To study the impact of matrix mechanical properties on pressure-driven dilation, PCT and mIMCD-3 cells were cultured under flow with intraluminal pressure in a denser 9 mg/mL collagen I matrix instead of 6 mg/mL Here representative PCT **(A)** and mIMCD-3 **(B)** tubules are shown in this configuration. On the right of these images are plotted the mean diameters of the tubules 5 days after confluency normalized by their mean diameters at the day of confluency (left, D5/D0) and 1 day after confluency (right, D5/D1). As reference, median and 95% confidence interval of analogous experiment in a 6mg/mL collagen gel are indicated by dark grey superimposed lines. **(A)** PCT tubules still display higher levels of dilation for *Pkd1^-/-^* cells (green, n=26) than parental ones (blue, n=28). **(B)** On contrary, for mIMCD-3 tubes there is no longer significant difference between the *Pkd1^-/-^* (green, n=25) and parental cells (blue, n=24). Boxplots shown are 1^st^ quartile, median and 3rd quartile, with whiskers showing extrema. Each dot represents a tubule (ns: p-value>0.05, *: p- value≤0.05, ****: p-value≤0.0001, actual p-values are indicated below the symbols). Scale bars: 100µm.

This matrix modification limited tubular dilation under flow + pressure conditions for parental cells of both cell types, but more markedly for mIMCD-3 cells. Dilations at day 5 fell for PCT at 46-68%, compared to 49-79% at 6 mg/mL, and for mIMCD-3 cells at 43-71%, compared to 78-110% at 6 mg/mL (95% CI) (Figure 3.A-B left, figure 2). Like for collagen 6 mg/mL, between day 1 and day 5, parental PCT tubules dilated by 20% on average when parental mIMCD-3 tubules exhibited levels of dilation way below 10%. The differences between PCT and mIMCD-3 cell lines were even more striking in the case of *Pkd1*^-/-^ cells. No decrease in dilation between tubules in a 6 and 9 mg/mL collagen matrix could be observed for *Pkd1*^-/-^ PCT, with diameter still increasing by 92- 128% at day 5 (Figure 3.A). However, *Pkd1*^-/-^ mIMCD-3 cells displayed a behavior much more sensitive to the matrix properties. Not only dilation decreased noticeably compared to tubules in 6mg/mL, but the tubule diameter increases at day 5 (41-57%) was no longer significantly different to parental tubules. Past the initial dilation, the mean tubule diameter appeared to stall, increasing only around 12% between day 1 and day 5 (Figure 3.B).

### 4. Proliferation and morphology of the epithelium upon hydrodynamic stimulations suggest the existence of two dilation mechanisms due to loss of *Pkd1*

The observed differences in tubular dilations as a function of hydrodynamic cues could be explained with differences either in net cell proliferation or/and with changes of cell morphology modulating the surface covered by the tissue. To unravel the cellular mechanisms involved in tubular dilation, we assessed levels of proliferation (KI67 labelling), as well as cell morphology. The mean internuclear distance within a tubule was used as a proxy for cell spreading.

In PCT tubules subjected to flow only (Figure 4.A), the levels of proliferation at day 1 were similar on average between parental and *Pkd1*^-/-^ cells (Figure 4.B). At day 5, most *Pkd1*^-/-^ tubules were still at least as proliferative as at day 1, with 5-9% Ki67 positive cells at day 1 and 9-16% (95%CI) at day 5 and were on average an order of magnitude more proliferative than parental cells tubules (Figure 4.B). Parental and *Pkd1*^-/-^ cells were hard to tell apart in terms of morphology and organization within the tissue at day 1, with no significant difference in internuclear distance. Sustained proliferation levels in *Pkd1*^-/-^ tubules resulted in denser tissues at day 5 as shown by cell morphology (Figure 4.A, S4-A) and was confirmed by a decrease in internuclear distance (Figure 4.C). Furthermore, in most instances we observed the formation of multilayers constituted of a few cells.

**Figure 4:**
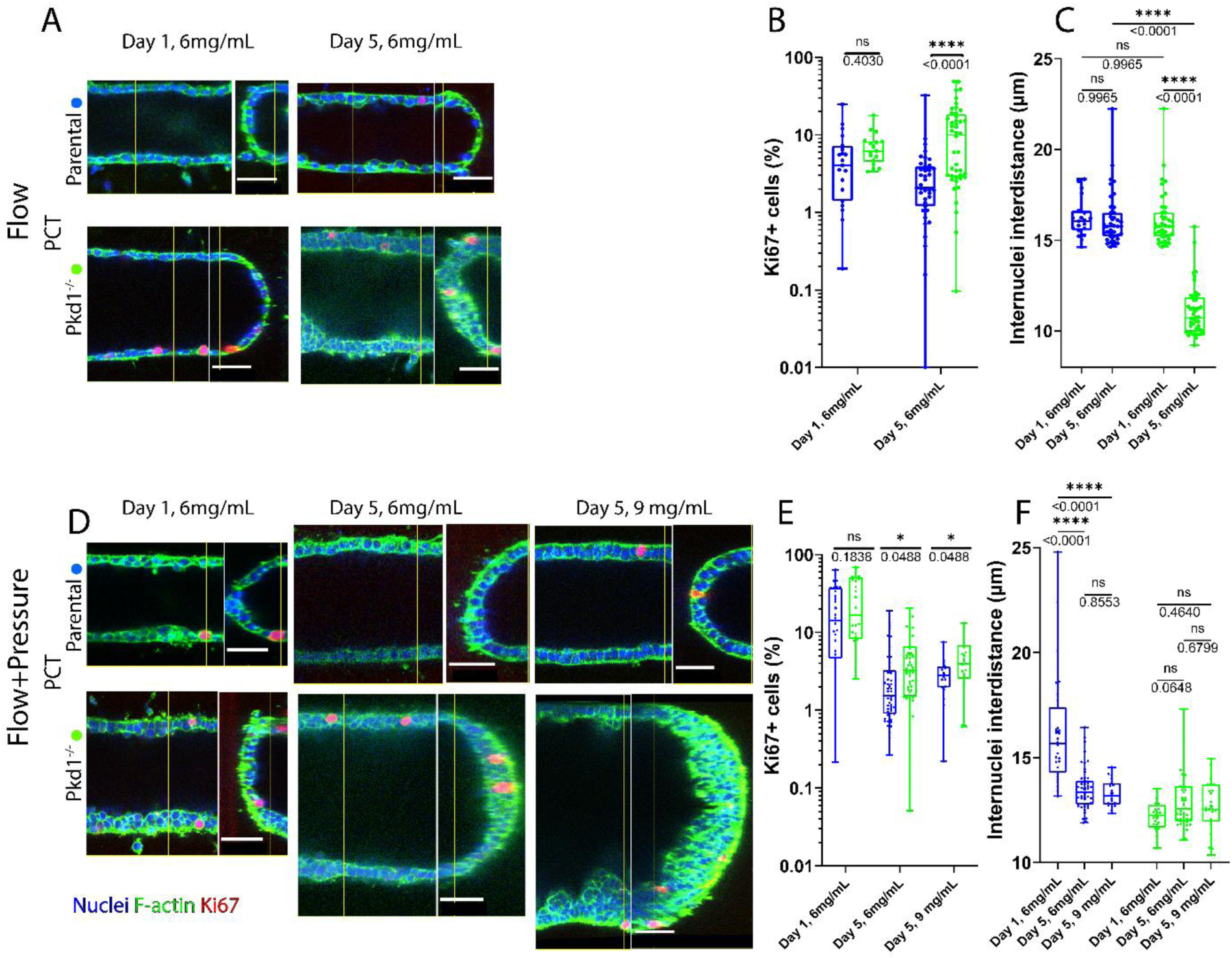
In PCT tubules, loss of *Pkd1* leads to higher proliferation levels and denser tissues. PCT tubules left under flow without pressure **(A)** or under flow with pressure **(D)** were fixed 1 day and 5 days after confluency in 6 mg/mL collagen I matrix, 5 days after confluency in 9 mg/mL collagen, and stained for nuclei (blue), F-actin (green), and Ki-67 (red). Cell proliferation was quantified by computing the ratio of Ki67 positive nuclei over the total number of nuclei **(B, E)**; comparisons between parental (blue) and *Pkd1^-/-^* cells (green) are shown. Cell morphology was assessed by measuring the average internuclear distance within each tubule **(C, F)** as a proxy for cell spreading. For internuclear distance, either parental (blue) or *Pkd1^-/-^* (green) values are shown side to side, to focus on the relative evolution in cell spreading related to time or matrix properties. **(B-C)** PCT tubes under flow, control *Pkd1^+/-^* at day 1 and day 5 (n=19 and n=39), and *Pkd1^-/-^* at day 1 and day 5 (n=17 and n=44), **(E-F)** PCT tubes under flow+pressure, control *Pkd1^+/-^* at day 1, day 5 and day 5 in 9mg/mL collagen (n=28, n=51 and n=19 respectively) and *Pkd1^-/-^* at day 1, day 5 and day 5 in 9mg/mL collagen (n=24, n=57 and n=23 respectively). Boxplots shown are 1^st^ quartile, median and 3rd quartile, with whiskers showing extrema. Each dot represents a tubule, (ns: p- value>0.05, *: p-value≤0.05, ****: p-value≤0.0001, actual p-values are indicated below the symbols). Scale bars: 50µm. Overall, *Pkd1^-/-^* cells exhibit higher proliferation levels than the parental cells. This correlates with *Pkd1^-/-^* tubules that become denser over time under flow and often form multilayer in all conditions.

Applying intraluminal pressure initially increased PCT proliferation, irrespective of *Pkd1* genotype (Figure 4.D). Overall, however, proliferation levels dropped drastically 5 days after confluency compared to experiments without intraluminal pressure. Nonetheless, *Pkd1*^-/-^ tubules sustained higher proliferation levels than parental cells tubules, with a statistically significant difference of 4-7% *vs*. 2-4% Ki67- positive cells respectively (Figure 4.E). This result suggests that for these cells intraluminal pressure initially induces a high rate of proliferation, irrespective of *Pkd1*, but importantly, over the course of 5 days, *Pkd1*^-/-^ cells are more proliferative than parental cells. Interestingly, this time, parental cells tubules became denser over time (Figure 4.F, S4.B). This coincides with these cells adopting a more cuboidal shape than with flow only, more reminiscent of *in vivo* proximal tubule cells. *Pkd1*^-/-^ cell morphology and density remained stable over time and comparable to parental cells. However, *Pkd1*^-/-^ cells still formed multilayers of a few cells. Increasing matrix density had no sizeable impact either on cell density or on proliferation (Figure 4E-F, S4.B).

In mIMCD-3 tubules with flow only (Figure 5.A), we observed a similar increase in proliferation at late stages (day 5) linked to the loss of *Pkd1*. Initially at day1, the proportion of Ki-67 positive cells was 9-16% and 10-27% (95%CI) for *Pkd1*^-/-^ and parental cells respectively, but it strongly increased to 21-29% for *Pkd1*^-/-^ cells at day 5 compared to only 3-6% for parental cells (Figure 5.B). This increased proliferation also translated into an increased density for mIMCD-3 *Pkd1*^-/-^ cells, shown by a decrease in internuclear distance (Figure 5.C). Contrary to PCT cells, this increased proliferation did not induce significant tubular dilation. It is worth noting that flow shear stress promoted for these cells a more cuboidal shape over time compared to our previous observations (Figure 5.A, S5-A).

**Figure 5:**
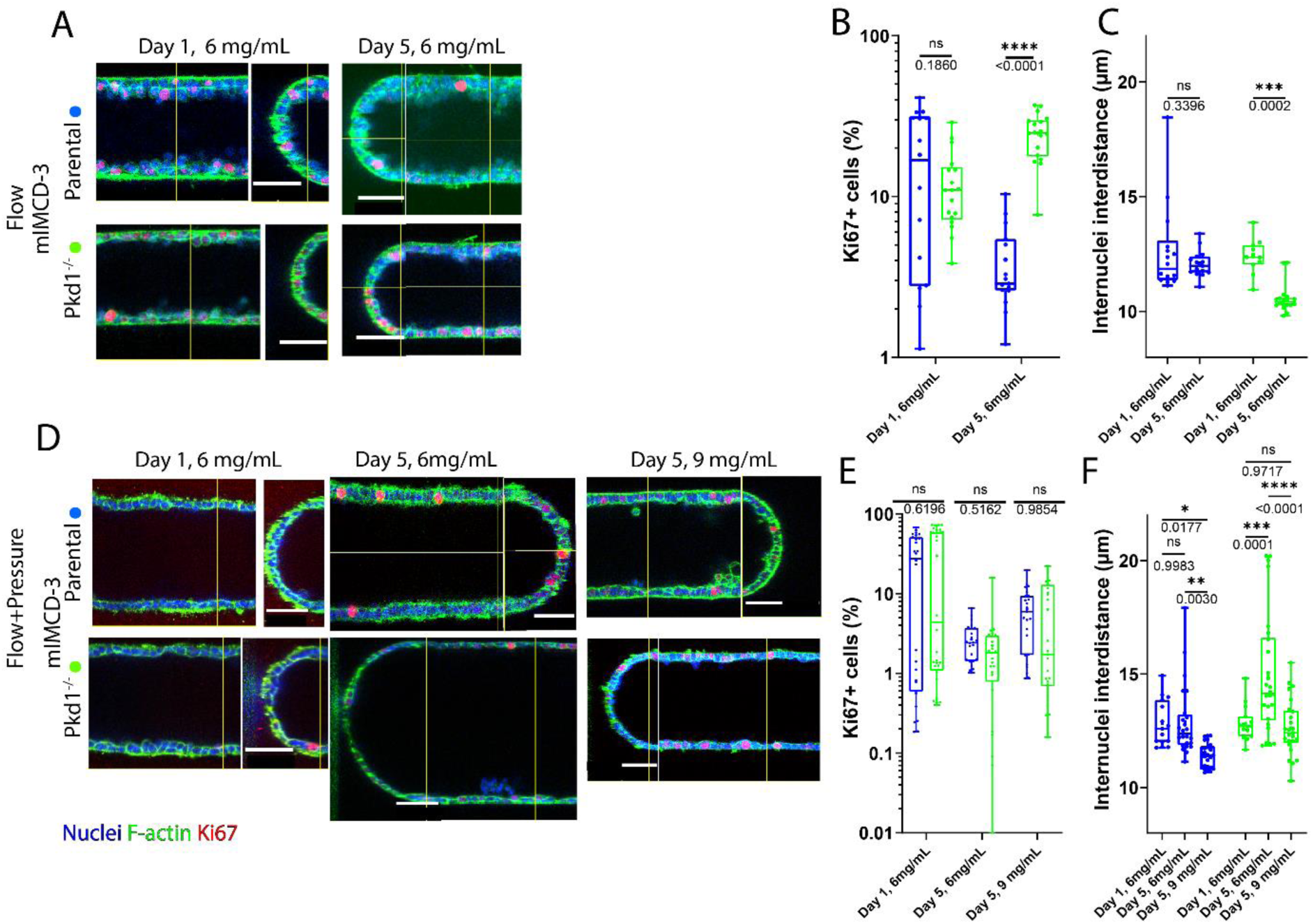
In mIMCD-3 tubules, increased tubular dilation is not correlated to higher proliferation levels but to increased cell spreading. mIMCD-3 tubules left under flow without pressure **(A)** or under flow with pressure **(D)** were fixed 1 day and 5 days after confluency in 6 mg/mL collagen I matrix, 5 days after confluency in 9 mg/mL collagen, and stained for nuclei (blue), F-actin (green), and Ki-67 (red). Cell proliferation was quantified by computing the ratio of Ki67 positive nuclei over the total number of nuclei **(B, E)**; comparisons between parental (blue) and *Pkd1^-/-^* cells (green) are shown. Cell morphology was assessed by measuring the average internuclear distance within each tubule **(C, F)** as a proxy for cell spreading. For internuclear distance, either parental (blue) or *Pkd1^-/-^* (green) values are shown side to side, to focus on the relative evolution in cell spreading related to time or matrix properties. **(B-C)** mIMCD-3 tubes under flow, control parental cells at day 1 and day 5 (n=14 and n=15), and *Pkd1^-/-^* cells at day 1 and day 5 (n=11 and n=18), **(E-F)** mIMCD-3 tubes under flow+pressure, control parental cells at day 1, day 5 and day 5 in 9mg/mL collagen (n=15, n=24, n=25 respectively) and *Pkd1^-/-^* cells at day 1, day 5 and day 5 in 9mg/mL collagen (n=15, n=26, n=25 respectively). Boxplots shown are 1^st^ quartile, median and 3rd quartile, with whiskers showing extrema. Each dot represents a tubule (ns: p- value>0.05, *: p-value≤0.05, **: p-value≤0.01, **: p-value≤0.001 ***: p-value≤0.0001, actual p-values are indicated below the symbols). Scale bars: 50µm. Contrary to PCT tubules, higher proliferation levels are associated with *Pkd1^-/-^* tubules only under flow without intraluminal pressure that become denser over time. Under flow with luminal pressure, however, if parental cells get significantly denser over time, *Pkd1^-/-^* cells become significantly more spread. This intraluminal pressure effect is mitigated by a denser matrix.

Contrary to PCT, addition of intraluminal pressure (Figure 5.D) did not initially have a sizeable impact on mIMCD3 proliferation (Figure 5.E). Interestingly however, at day 5 the ratio of *Pkd1*^-/-^ cells in the cell cycle appeared to have decreased by at least an order of magnitude and there was no longer a significant difference with parental cells. In contrast, the difference in cell morphology and density between *Pkd1*^- /-^ and parental cells was striking (Figure 5.D, S5-B). Initially it was hard to distinguish between *Pkd1*^-/-^ and parental cells based on appearance and by using the internuclear distance criteria (Figure 5D-E, S5-B, day1). However, at day 5 *Pkd1*^-/-^ cells adopted a squamous morphology and the tissue became less dense (Figure 5D-E, S5-B, day5). Parental cells followed an opposite trend wherein the cells looked more cuboidal keeping a similar cell density. These results suggest that, for mIMCD-3, the excessive dilation of *Pkd1* tubule does not originate from enhanced proliferation but is correlated with the acquisition of a squamous morphology in response to intraluminal pressure. Increasing the matrix concentration favored the formation of denser tissues in both parental and *Pkd1^-/-^* mIMCD-3 tubes (Figure 5D-E, S5-B, day5, 9mg/mL). For *Pkd1*^-/-^ cells in particular, the cell morphology was similar to the initial morphology in a 6mg/mL matrix, which further suggests that *Pkd1*-dependent tubular dilation for mIMCD-3 and an abnormally squamous cell morphology are two intertwined events.

## Discussion

In a kidney-on-chip with deformable tubes of physiological dimensions, formed in a scaffold with tunable mechanical properties, we developed a method to decouple flow and luminal pressure. We propose here an innovative solution to decouple two generally intricate mechanical stimuli, pressure, and flow shear stress, relying on classic microfabrication technics and perfusion with ubiquitous material. Our platform is modulable and allows to run in parallel a great number of different replicates for different mechanical conditions. We applied this system to specifically investigate how these mechanical parameters affect the shape and organization of tubules formed by kidney epithelial cell lines derived either from proximal (PCT) or distal (mIMCD-3) mouse nephron. In each of the cell lines, we further studied how these mechanical parameters affect the excessive tubule dilation caused by polycystin-1 inactivation.

Compared to the results that we obtained in static conditions (Myram et al^44^., Figure S2, Mazloum et al^33^), applying flow shear stress with negligible pressure helped to maintain a stable hollow tubular geometry, limiting epithelial multilayering and cell accumulation in tubule lumen. Irrespective of the cell line considered, further applying pressure led to a progressive increase in tubule diameter, suggesting that all the studied cell lines respond to pressure. This pressure-induced dilation appeared to be initially independent of genotype and can be most likely attributed to the mechanical stimuli induced by the elastic response of the confluent tubules. However, this initial response appeared to be followed by a cell-specific deformation of the tubules, indicating that different mechanisms more complex than a response to an elastic deformation under pressure are at play. Overall, cell responses to these mechanical parameters were contrasted, depending not only on *Pkd1* status but also on the cell line considered. PCT expressing *Pkd1* adopted a more physiological shape when exposed to pressure, becoming more cuboidal and forming denser tissues only when adding intraluminal pressure, a behavior not observed in mIMCD-3 cells. This differential response may be linked to the physiologic exposure to pressure of the nephron segments from which those cell lines originate, as pressure drops from the proximal tubule to the distal nephron.

Strikingly, while *Pkd1* deletion in both PCT and mIMCD-3 led to excessive tubule dilation in static conditions, this phenotype was differentially affected by mechanical parameters across cell lines. Essentially, *Pkd1-*deficient PCT tubules underwent excessive dilation, increased proliferation and cell crowding, irrespective of pressure, flow or ECM stiffness conditions. This excessive proliferation of *Pkd1*^-/-^ PCT further led to cell crowding and epithelial multilayering, suggesting defective contact inhibition.

In contrast, the ability of *Pkd1*-deficient mIMCD-3 cells to deform their scaffold was drastically impacted by the mechanical conditions applied. *Pkd1*^-/-^ mIMCD-3 cells dilated the ECM scaffold more than the parental line in static conditions in a 6 mg/mL collagen scaffold (Figure S2, Mazloum et al^33^). Applying shear stress alone prevented the excessive dilation of the tubule formed by *Pkd1*^-/-^ mIMCD-3 despite an increased proliferation. The absence of multilayering may suggest an enhanced regulation of the tissue homeostasis possibly through apical extrusion. Surprisingly, adding luminal pressure overcame the protective effect of flow shear stress and restored the ability of *Pkd1*^-/-^ to distend the ECM scaffold. In contrast, wild-type mIMCD-3 cells lacked in most cases this ability to distend the ECM past a rapid initial dilation. Increasing ECM concentration prevented the excessive dilation of the tubules formed by *Pkd1*^-/-^ mIMCD-3 under flow+pressure conditions. In addition, contrary to PCT cells, the excessive dilation of the tubule formed by *Pkd1*-deficient mIMCD-3 under pressure was not caused by increased cell proliferation but was associated with cell flattening, which was also prevented by an increase in ECM concentration. This suggests that cell flattening and tubule dilation for these cells are intertwined events in a tubule distension.

Interestingly, a discrepancy between the effect of ECM concentration over this distension and the associated matrix stiffening points to a nonlinear response of the tubule pertaining to an increase in matrix concentration. This nonlinear relationship between tissue mechanics and tubule response highlights how -cell matrix interactions play a critical role in shaping *Pkd1^-/-^* mICD3 tubule. Note that the basement membrane deposited by cells was not directly quantified here; nonetheless, *in vivo* parallel experiments demonstrated that early distal dilation were accompanied by a thinning of the basement membrane and the secretion of matrix softeners^33^. Furthermore, it has already been observed that loss of *Pkd1* in other *in vitro* models with mIMCD-3 cells is accompanied by abnormal matrix deposition and cell-matrix interactions^54^.

Our results unveil two distinct mechanisms leading to tubular dilation due to the loss of *Pkd1*. For the PCT, over-proliferation mechanically leads to tubular dilation regardless of mechanical constraints, hydrodynamic or from the extracellular matrix properties. For mIMCD-3 cells, a squamous cell phenotype favors tubular dilation through a mechanism both sensitive to the matrix properties and intraluminal hydrodynamic constraints but not associated with increased proliferation. The surprising results obtained with mIMCD-3 remarkably recapitulates our parallel *in-vivo* results^33^ regarding the initial dilation of *Pkd1*-deficient distal tubule mice, which undergoes distension without a detectable increase in tubular cell proliferation index. The effect of pressure on tubule-on-chip dilation further echoes the effect of ureteral obstruction, which precipitates cyst formation specifically in the distended distal nephron. Importantly, the finding that the ability of *Pkd1*- deficient mIMCD-3 cells to dilate tube is not only impacted by flow shear stress but also by luminal pressure and matrix stiffness suggests that PC1 is connected to different mechanosensitive circuits responding to distinct physical parameters.

The results obtained with PCT only partially coincide with the behavior of *Pkd1*- deficient proximal tubule *in vivo*. Increased cell proliferation and relative insensitivity to increased pressure are notable similarities. However, cell crowding as well as the striking formation of a multilayered epithelial layers are not observed in most orthologous ADPKD mice models^33,55^. This phenotype is nonetheless reminiscent of the micro-polyp and cord-like hyperplasia features found in a few percent of human ADPKD cysts^56^.

In conclusion, with this work, we show the importance of a specific control of intraluminal pressure in kidney tubule models, more particularly in the context of polycystic kidney disease. Here, it allowed us to demonstrate that even moderate levels of hydrostatic pressures have specific effects on kidney cells in pathological and non-pathological contexts, with mechanisms that may differ depending on the nephron segment. Although there were recent studies in different systems (blood microvessels^57^) decoupling pressure from flow shear stress, tubular kidney models have not separately dissected the role of these two mechanical stresses arising from perfusion. Thus, with future *in vitro* kidney models, a specific control of the effects of flow and pressure could be pivotal to decipher the mechanisms underlying diseases such as ADPKD but also more generally, to improve the biomimicry of these systems.

## Acknowledgements.

The authors acknowledge all the members of the MMBM and PBME groups for their precious support in the process development and in microfabrication. We are grateful to Sarah Myram, Sophie Demolombe, Eric Honoré, Bénédicte Delaval, Abdul Barakat, Jacques Fattacciolo, Pierre Ucla, Isabelle Bonnet, Pascal Silberzan and Axel Buguin for insightful discussions, and to Polina Rapoport and Lorraine Couteau for their participation in the experiments. We thank George M. O’Brien Kidney Center at Yale (NIH P30 DK079310) and Stefan Somlo for the kind gift of PCT *Pkd1^+/-^* and PCT *Pkd1^-/-^* cell lines and Michael Köttgen, Lukas Westermann and Tilman Busch, Freiburg University, for the kind gift of mIMCD-3 control and *Pkd1*-KO cell lines. This work benefited from the technical contribution of the joint service unit CNRS UAR 3750. The authors would like to thank the engineers of this unit (in particular Bertrand Cinquin) for their advice during the development of the experiments. The authors greatly acknowledge the Cell and Tissue Imaging core facility (PICT-IBiSA), Institut Curie, member of the French National Research Infrastructure France-BioImaging (ANR10-INBS-04).

## Author Contributions

B.L., F.B., S.D. and S.C. designed the study; B.L., G.G.,

J.V. and M.M. performed experiments; B.L., J.V., S.D. and S.C. analyzed data; S.D. and S.C. supervised the study and wrote the manuscript; B.L. S.D. and S.C. designed the study, drafted, and revised the manuscript; all authors approved the final version of the manuscript.

