## Supplemental Information for "Decoupling shear stress and pressure effects in the biomechanics of autosomal dominant polycystic kidney disease using a perfused kidney-on-chip"

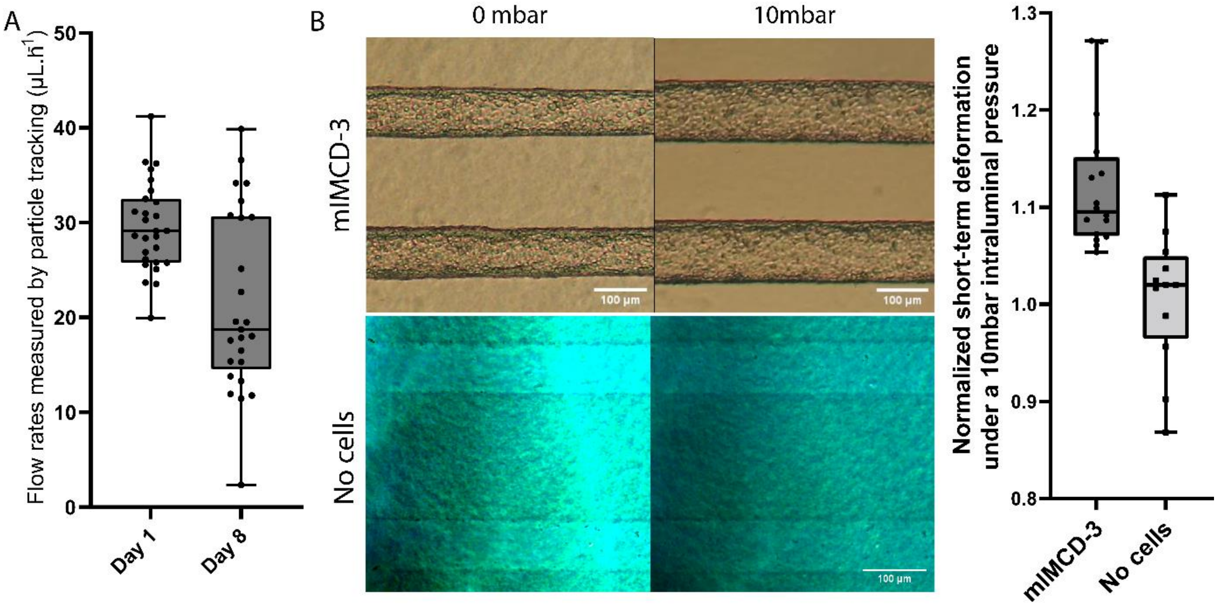


**Figure S1: Measured flow rates in perfused chips seeded with mIMCD-3 cells and tubular viscoelastic dilation under intraluminal pressure with or without cells.**

We measured flow rates in tubules lined with mIMCD-3 cells 1 day and 8 days after confluency (A) as well as their initial elastic dilation right after confluency due to a 10mbar pressure (n=16) and compared them to the response of naked collagen tubules (n=12) (B). To do so, we measured the diameter of the tubule perfused in the flow+pressure configuration a few minutes after applying a 10mbar pressure and normalized by their diameter right before this perfusion. Boxplots shown are 1$^{st}$

quartile, median and 3rd quartile, with whiskers showing extrema. Each dot represents a tubule.

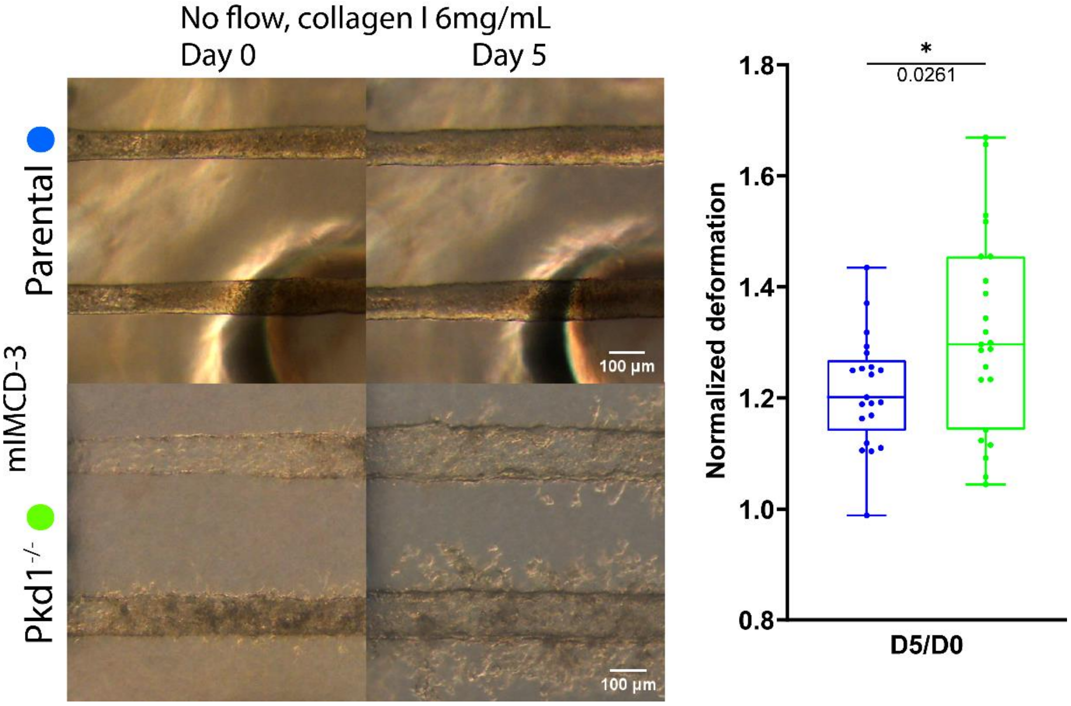


**Figure S2: Static dilation of mIMCD-3 tubules depending on *Pkd1*.**

Tubules were seeded with parental (n=23) and *Pkd1*$^{-/-}$ (n=21) mIMCD-3 cells and maintained without perfusion. Here are shown representative tubules the day of confluency and at five days of confluency. On the right are plotted their mean diameters five days after confluency normalized by their diameter the day of confluency. Boxplots shown are 1$^{st}$ quartile, median and 3rd quartile, with whiskers showing extrema. Each dot represents a tubule (ns: p-value>0.05, *: p-value≤0.05, ****: p-value≤0.0001, actual p-values are indicated below the symbols).

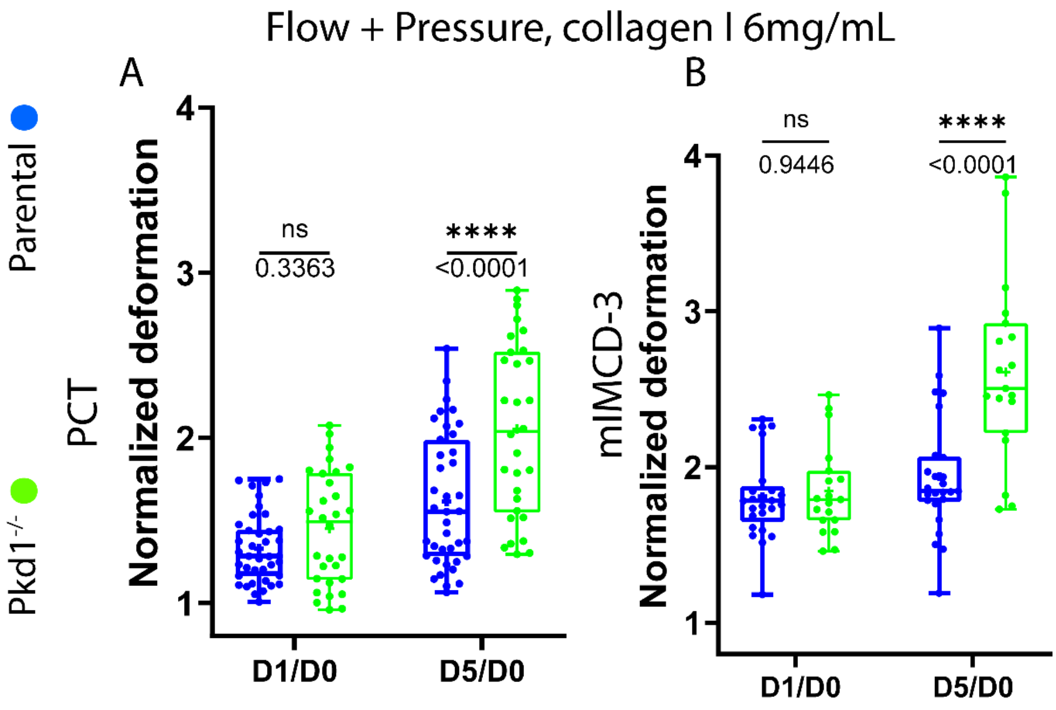


**Figure S3: Normalized dilation of perfused PCT and mIMCD-3 tubules under flow and pressure.**

Here are plotted ratios showing the temporal evolution of the mean diameters of parental (blue) and *Pkd1*$^{-/-}$ (green) PCT (A, n=39 and 30 respectively) and mIMCD-3 (B, n=25 and 19 respectively), 1 day and 5 days after confluency normalized by their diameters at confluency (day 0). Ratios D1/D0 are shown (left), and ratios D5/D0 already shown on Fig. 2C,D are shown here again for comparison to the evolution at larger timescales. Boxplots shown are 1$^{st}$ quartile, median and 3rd quartile, with whiskers showing extrema. Each dot represents a tubule (ns: p-value>0.05, *: p-value≤0.05, ****: p-value≤0.0001, actual p-values are indicated below the symbols), statistical tests ANOVA.

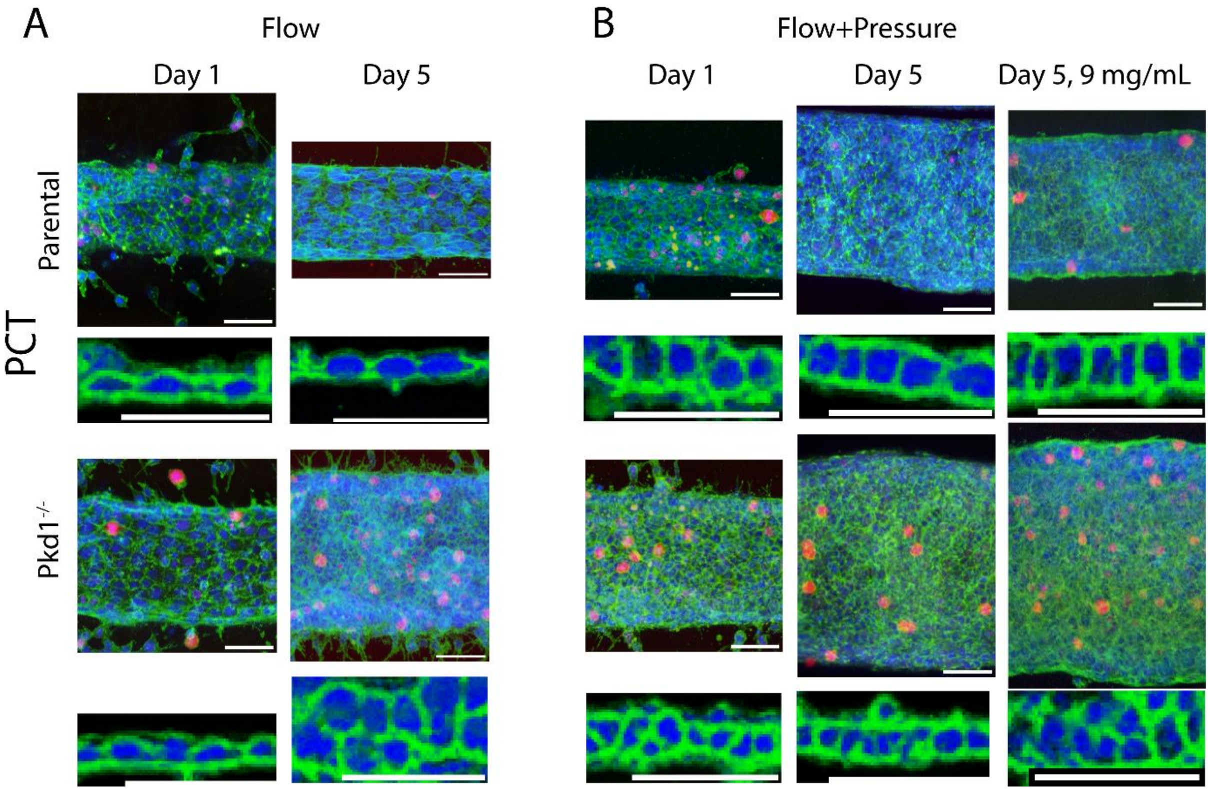


**Figure S4: Confocal images of representative PCT tubules and zoom on corresponding pre-segmented cells 1 day or 5 days after confluency.** Fluorescence: blue: nuclei, green: actin, red: Ki-67. Pairs of images, with *z* projections on top and representative side sections on the bottom. Scale bar: 50µm.

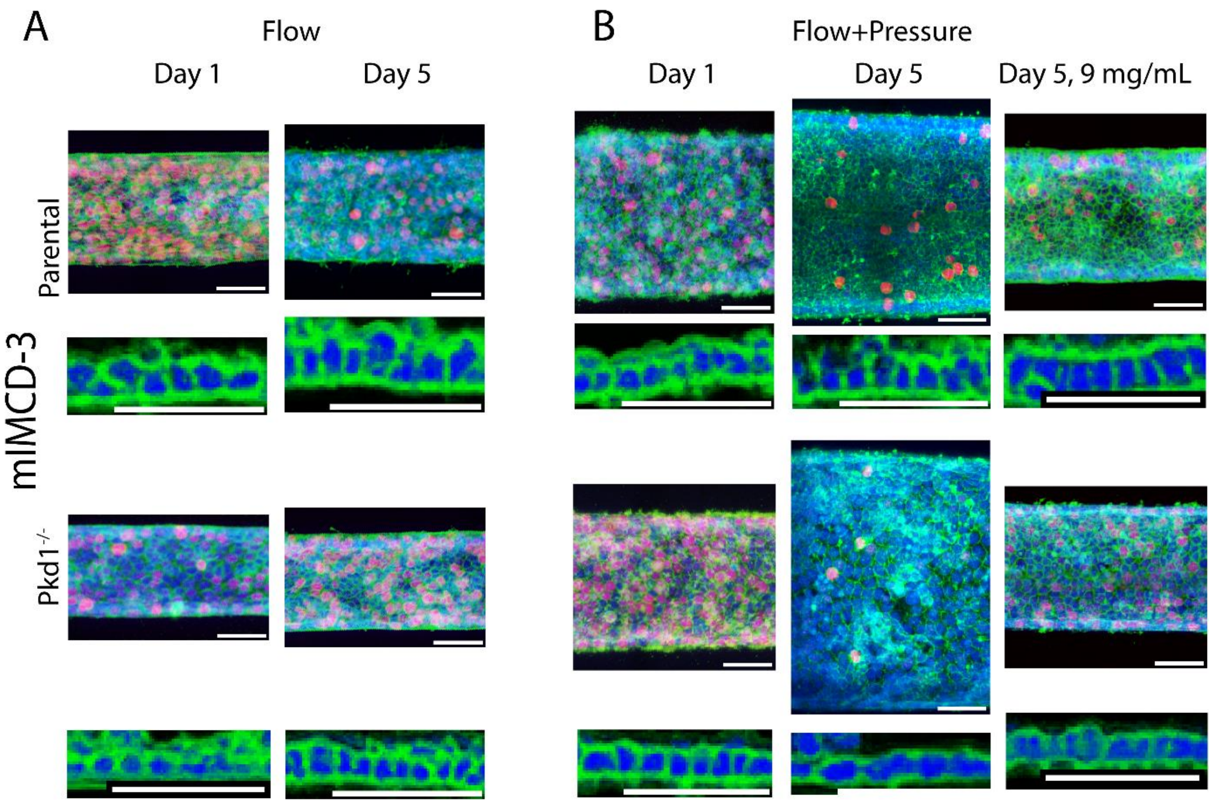


**Figure S5: Confocal images of representative mIMCD-3 tubules and zoom on corresponding pre-segmented cells 1 day or 5 days after confluency.** Fluorescence: blue: nuclei, green: actin, red: Ki-67. Pairs of images, with *z* projections on top and representative side sections on the bottom. Scale bar: 50μm.
